# DNA spontaneously wrapping around a histone core prefers negative supercoiling: A Brownian dynamics study

**DOI:** 10.1101/2024.08.05.606726

**Authors:** Chunhong Long, Hongqiong Liang, Biao Wan

## Abstract

In eukaryotes, DNA achieves a highly compact structure primarily due to its winding around the histone cores. The nature wrapping of DNA around histone core form a 1.7 left-handed superhelical turns, contributing to negative supercoiling in chromatin. On the contrary, negative supercoils generated behind the polymerase during transcription may play a role in triggering nucleosome reassembly. To elucidate how supercoils influence the dynamics of wrapping of DNA around the histone cores, we developed a novel model to simulate the intricate interplay between DNA and histone. Our simulations revealed that both positively and negatively supercoiled DNAs are capable of wrapping around histone cores to adopt the nucleosome conformation. Most of all, our findings confirmed a preference for negative supercoiled DNA during nucleosome wrapping, and revealed that the both of the negative writhe and twist are comparatively beneficial to the formation of the DNA wrapping around histone. This advancement in the understanding of spontaneously nucleosome formation may provide insights into the intricate dynamics of chromatin assembly and its diverse functions. Our model thus can be further utilized to simulate the formations of multi-nucleosomes during re-assembling of the chromatin fiber, which will significantly enhance the understanding of the fundamental mechanisms governing the structure and function of chromatin.

**Author summary:** The compacted organization of DNA within chromatin is primarily attributed to its intricate winding around histone cores. This winding process involves 1.7 left-handed superhelical turns around the histone core, contributing to the negative supercoiling within chromatin fibers. To gain deeper insights into how DNA supercoiling impacts the dynamics of DNA wrapping around histone cores, we devised a novel computational model that simulates the intricate interplay between DNA and histone cores. Additionally, our simulations revealed that both of the positively and negatively supercoiled DNA can spontaneously adopt the nucleosome conformation upon wrapping around the histone core, and demonstrated a preferential tendency for negative supercoiling during nucleosome wrapping. Finally, we examined that both the negative writhe and twist components are comparatively advantageous for the formation of the nucleosome. The studies shed light on the intricate dynamics underlying chromatin assembly and its functional implications.

## Introduction

In eukaryotes, the primary step in the intricate folding of DNA into chromatin involves the wrapping of DNA strands around the histone core [1]. The crystal structure of the histone core particle of chromatin contains two copies of each histone protein, H2A, H2B, H3 and H4 being assembled into an octamer that accommodates approximately about 145 base pairs (bp) wrapped around it [2–5]. The negatively charged DNA wraps around the positively charged histone proteins about two turns in a counter-clockwise fashion [6, 7]. The left-handed wrapping of DNA around the histone octamer results in the formation of approximately 1.7 superhelical turns [1, 8]. The packaging of DNA within a robust nucleosome framework in eukaryotes physically obstructs the establishment of the initial replication and transcription complexes, as well as the subsequent activities of the respective machineries. Conversely, during transcription elongation, the positive supercoils generated in front of the RNA polymerase may destabilize the nucleosome while the negative supercoils behind the polymerase may aid in triggering the nucleosome reassembling [9, 10]. Recent biochemical and molecular biological experiments have focused on stretching DNA molecules to investigate their mechanical or dynamical behaviors [6, 11–14]. Meanwhile, some theoretical works have also explored the mechanical or dynamical properties of DNA and chromatin [15–17]. Along with experimental and theoretical studies of DNA and chromatin, numerous molecular dynamics simulation studies had been done on structures and stabilities of nucleosomes [18–25]. Recently, the Brownian dynamics methods had been developed, which have been employed to study the kinetics of DNA and nucleosome interactions, providing further insights into the dynamic nature of chromatin assembly and function [15, 26–32].

In this study, we developed a Brownian dynamics model to investigate the dynamics properties of spontaneously nucleosome wrapping by supercoiled DNA. Our model integrated a discrete worm-like chain and a rotational sphere, for representing DNA and histone, respectively. Employing this model, we confirmed that the interaction between histone and DNA is the primary driving force behind nucleosome wrapping, regardless of the supercoiling density of the DNA. Furthermore, our simulations demonstrated that negative supercoils are more beneficial to spontaneous nucleosome wrapping, which supports that negative supercoils facilitate the wrapping of DNA around histone octamers, thereby promoting the formation of nucleosomes. And finally we discussed the mechanisms underlying the preference for negative supercoiling during nucleosome wrapping. Our findings provide insights into how the DNA supercoiling impacts chromatin structure, function and assembly.

## Method

Our coarse grained model consists of a discrete worm-like chain and a rotational sphere, depicted in FIG 1, to simulate the supercoiled DNA and the histone core, respectively.

**Fig 1.**
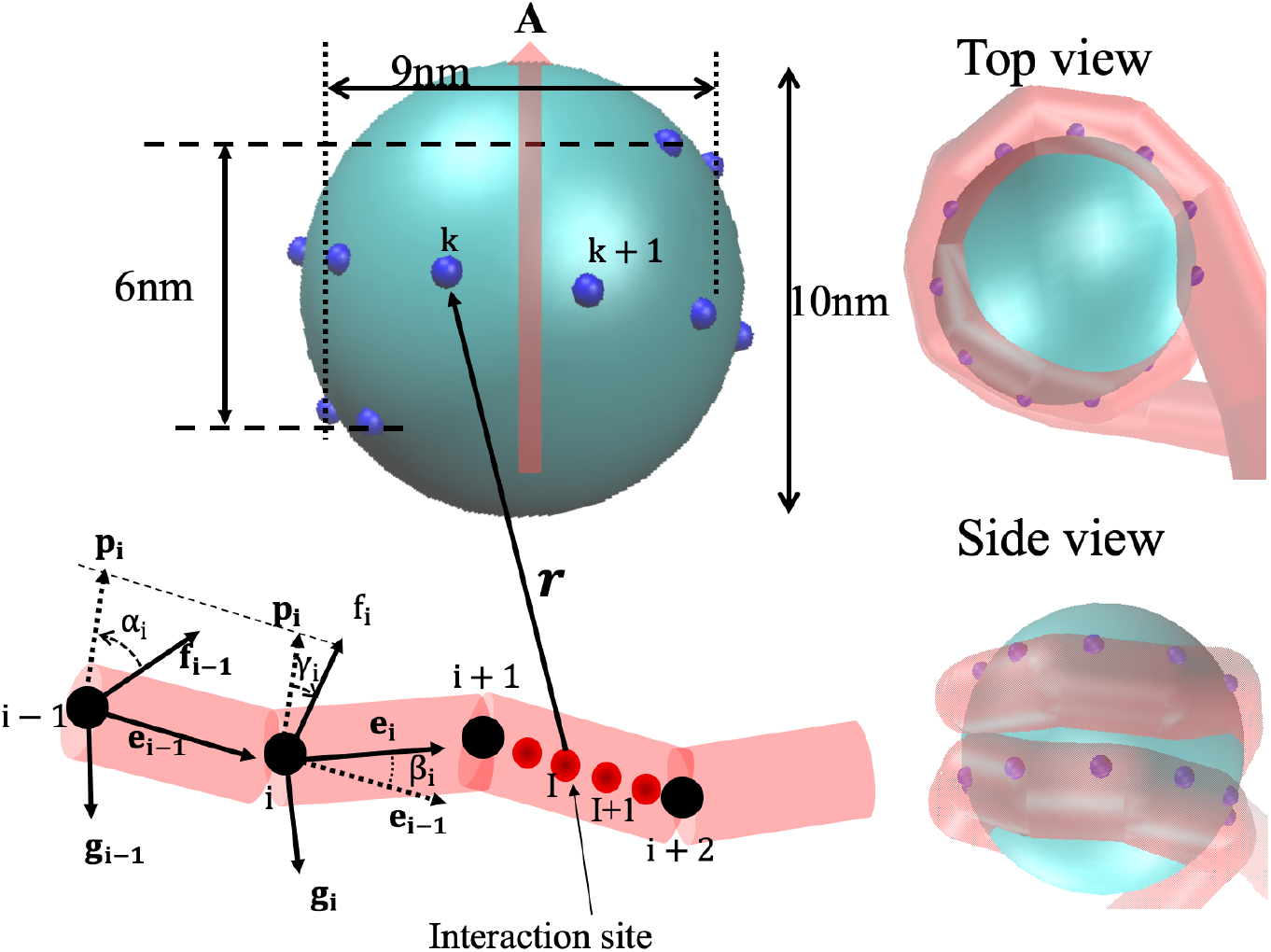
Schematic view of the histone core and DNA interaction model. The histone core is modeled as a sphere entity, possessing 20 interaction points along a left-handed helical path that enable the DNA to be wound around the histone core. The axial vector **A** of the helical path on the sphere describes the orientation of the histone core. The DNA is modeled as *N* discrete segements. The *i*-th segment is defined by vertices *i* and *i* + 1, attached by **f**_*i*_, **g**_*i*_ and **e**_*i*_. The vectors define the bending angle *β*_*i*_ = arccos(**e**_*i*−1_ · **e**_*i*_), and the twisting angle *θ*_*i*_ = *α*_*i*_ + *γ*_*i*_ by introducing an auxiliary vector **p**_**i**_ = **e**_**i**−**1**_ × **e**_**i**_. (Right) Two views of the nucleosome structure generated by VMD from the simulations.

The discrete worm-like chain is comprised of *N* jointed discrete segments (cylinders) [33]. The conformation of this chain is described by *N* + 1 vertices {**r**_*i*_}. The i-th segment is defined by the vector **s**_*i*_ ≡ **r**_*i*+1_ − **r**_*i*_. The equilibrium length of each segment is *l*_0_ = 3.4*nm*, representing 10 bp. In addition, an orthogonal base vector frame {**e**_*i*_, **f**_*i*_, **g**_*i*_} (**e**_*i*_ = **f**_*i*_ × **g**_*i*_) is attached to the segment (Fig 1), where **e**_**i**_ ≡ **s**_**i**_/*s*_*i*_, **f**_*i*_ and **g**_*i*_ describe a local rotational angle *ϕ*_*i*_ of the segment about its axis **e**_*i*_. The torsion of the chain is described by *N* rotational angles {*ϕ*_*i*_}. The i-th bending angle *β*_*i*_ is defined as arccos (**e**_*i*−1_ · **e**_*i*_). The i-th twist *θ*_*i*_ depends on the two frames {**f**_*i*−1_, **g**_*i*−1_, **e**_*i*−1_} and {**f**_*i*_, **g**_*i*_, **e**_*i*_}. By introducing an auxiliary vector **p**_**i**_ = **e**_**i**−**1**_ × **e**_**i**_, the twist is *θ*_*i*_ = *α*_*i*_ + *γ*_*i*_, where *α*_*i*_ is the rotation angle from **f**_*i*−1_ towards **p**_**i**_, *γ*_*i*_ is the angle from **p**_**i**_ towards **f**_*i*_.

The stretching, bending, twisting potential energies are all harmonic [33]. The electrostatic interaction is represented as the Debye-Hückel potential on the uniformly distributed point-like charges on the segments [34–36]. Thus the energetics of the DNA chain is

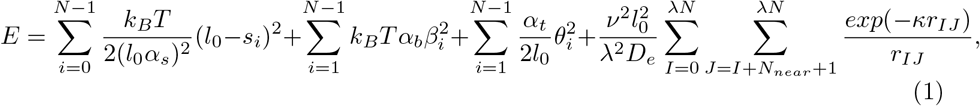

where the first term on the right-hand side is the stretching energy of each segment and *α*_*s*_ is a stiffness coefficient; the second term is the bending energy and *α*_*b*_ is a bending stiffness coefficient; the third one is the twisting energy and *α*_*t*_ is a torsional stiffness coefficient [33]; the last one is the electrostatics and *ν* and *λ* are the effective charge density on each segment and the number of the point-like charges per segment, respectively, *D*_*e*_ is the dielectric constant of water, *N*_*near*_ is the number of neighbor point-like charges of each segment, and *κ* is the inverse Debye length. Moreover, the impenetrability of DNA segments is guaranteed by introducing a repulsion force 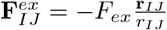 if *r*_*I,J*_ < 2*nm* [34], where the indices I and J denote the point-like charges, not the vertices. All the simulation parameters used in this method are listed in the TABLE 1.

**Table 1.** Constants and parameters.

| Parameter | Symbol | Value [33, 35] |
| --- | --- | --- |
| Number of segments | $N$ | 100 |
| Temperature | $T$ | 298K |
| Histone core radius | $R_h$ | 5nm |
| Segment equilibrium length | $l_0$ | 3.4nm |
| Stretching stiffness coefficient | $\alpha_s$ | 0.1 |
| Bending stiffness coefficient | $\alpha_b$ | 7.32 |
| Torsional stiffness coefficient | $\alpha_t$ | 400pN · nm<sup>2</sup> |
| Monovalent salt concentration |  | 150 mM |
| Dielectric constant | $D_e$ | $8.9 \times 10^{-9} F/m$ |
| Effective charge density | $\rho_e$ | 8.01e/nm |
| Inverse Debye length | $\kappa$ | 1.261nm<sup>-1</sup> |
| Number of the point charges per segment | $\lambda$ | 5 |
| Electrostatic cutoff distance | $r_c$ | 8nm |
| DNA excluded volume force | $F_{ex}$ | 15pN |
| Viscosity | $\eta$ | 0.001kg/(m · s) |
| Hydrodynamic DNA radius | $R_H$ | 1.3nm |
| Hydrodynamic DNA rotational radius | $R_d$ | 1.2nm |
| Simulation time-step size | $\Delta t$ | 25ps |

The histone core is considered as a rotational sphere with diameter 10*nm*, on which 20 discrete interaction points are distributed along a left-handed helical path, spanning approximately 1.7 turns. The helical path of these interaction sites extends vertically for a height of 6 nm, while the diameters of the helix’s apical and basal termini approximate 9 nm.

The interactions between the DNA and the points on the sphere are modeled as the Morse potential [29]

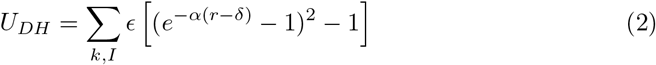

where *E* is the absorption energy, appropriately chosen as 5*k*_*B*_*T* (corresponding to the absorption energy density 8*k*_*B*_*Tnm*^−1^) [29, 30], *α* ∼ 1.26*nm*, is the inverse decay length, *δ* = 1*nm*, specifies the radius of the repulsive core of DNA, *r* is the distance between the *I*-th DNA binding site and the *k*-th site on the histone. The excluded volume effect between DNA and histone is considered by a half-harmonic potential, i.e., for *r* < |*a*| + *r*_0_,

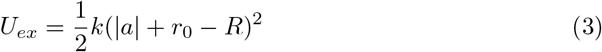

where *k* = 2*k*_*B*_*T* is the rigidity coefficient, |*a*| = 5*nm* is the radius of the histone, *r*_0_ = 1*nm* is the radius of DNA and *R* is the distance between the *I*-th DNA binding site and the center of the histone.

The Brownian dynamics (BD) can be performed using an integration, the second-order BD algorithm [33, 35]. A tentative first-order displacement is

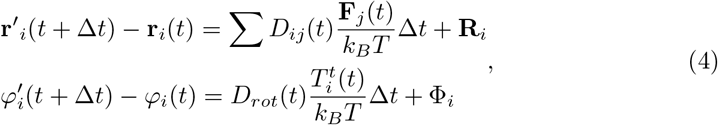

and the final half-step is

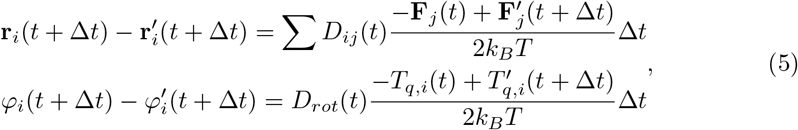

where 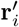 is the tentative position, *D*_*ij*_ is the Rotne-Prager tensor component [33], and for the computational efficiency, we chose *D*_*ij*_ = *Dδ*_*ij*_ = *δ*_*ij*_*k*_*B*_*T*/6*πηR*_*H*_, and here *R*_*H*_ is effective radius of each segment [35, 37], **F**_*i*_ and 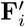 are the forces on the *i*-th vertex corresponding to the conformation **r** and **r**^*l*^, respectively, *T*_*q,i*_ and 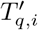 are the torques on the *i*-th segment; 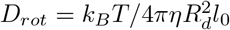 is the rotational diffusion constant, here *R*_*d*_ is the diffusion radius of DNA, **R** and Φ are the thermal noises with the properties (**R**) = 0, (**R** ⊗ **R**) = 2**D**Δ*t*, (**Φ**) = 0, (**Φ** ⊗ **Φ**) = 2**D**_**rot**_Δ*t*.

The center of mass of the histone protein moves following the Langevin equation under the protein-DNA interactions *U*_*DH*_ + *U*_*ex*_,

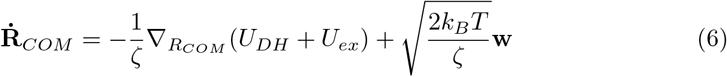

where *ζ* = 6*πη*|*a*| is the the friction coefficient, with *η* the solution viscosity and |*a*| the radius of the spherical protein, and *w*(*t*) is the Gaussian noise with zero mean and unit variance. It should be noted that the BD (EQ.6) can be performed using the second-order BD algorithm.

The orientational or rotational degrees of the histone core caused by thermal fluctuations and interactions are modeled by partitioning the rotations into the spatial rotation of the unit axial vector **A** (axial vector of the helical path), Δ**Ω**_*A*_ and the spinning about **A**, Ψ [38]. Accordingly, the protein angular Langevin dynamic equations are formulated as (details in Text S1).

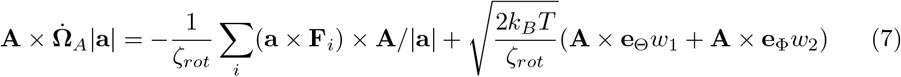

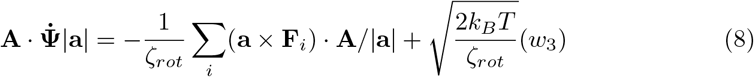

where *ζ*_*rot*_ = 8*πη*|*a*| is the friction due to rotation of the protein in solution, *w*_1_,*w*_2_ and *w*_3_ are the Gaussian noises with zero mean and unit variance. The source code is available at: https://figshare.com/s/b38438c966505126910e.

## Results

### Spontaneously wrapping of DNA around histone

A relaxed B-DNA consisting of *N* base pairs has a linking number *Lk*_0_ = *N*/10.5. The excess linking number difference of a supercoiled DNA is then Δ*Lk* ≡ *Lk* − *Lk*_0_. The supercoil density is accordingly defined as 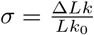. When a histone is completely wrapped by DNA, which contributes to approximately −1 linking number difference.

In FIG.2, an extended and supercoiled DNA (Δ*Lk* = 3, *i*.*e*.,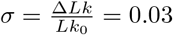) at length 1000*bp* stretched by *f* = 1.0*pN* interacts with a histone. The interactions between DNA and histone with absorption energy *E* = 5*k*_*B*_*T* can wind the DNA around the histone core, forming a partially wrapped structure, and finally trigger the completion of the nucleosome wrapping.

**Fig 2.**
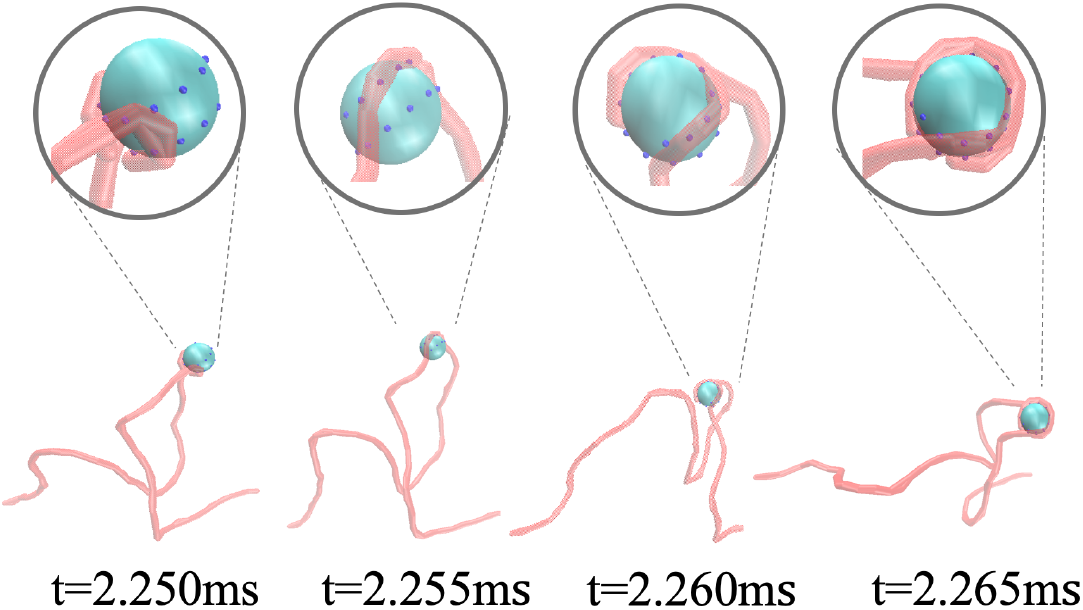
Representative conformations of DNA winding around a histone core obtained from the Brownian dynamics simulations. A single histone core interacts with a positively supercoiled DNA (*σ* = 0.03) at length of 1000 bp (100 segments) under tension *f* = 0.1*pN*.

It should be noted that the nucleosome state induces an extra ΔΔ*Lk* = +1 to the unconstrained DNA, which is energetically adverse. However, our simulation suggests that the positive supercoil is able to wrap a histone to form nucleosome.

### Strong interaction between histone and DNA promotes the nucleosome wrapping

The DNA winding around the histone core is energetically costly. Hence, the interactions between the DNA and histone should be the primary factor for nucleosome wrapping. The conformations exhibited by the histone-DNA complex can be broadly categorized into three types: mis-wrapped (non-left-handed), partially wrapped and nucleosome wrapped states(FIG.3(a)). The typical pathways of the nucleosome wrapping are *mis-wrapped state*→*partially wrapped state*→*nucleosome state* and *partially wrapped state*→*nucleosome state* (see the Text S2 for the details).

Intuitively, the strong interactions can promote the completion of the nucleosome wrapping. This is confirmed by our simulations of spontaneously nucleosome-wrapping with different DNA supercoil densities −0.05 ≤ *σ* ≤ 0.05 and different adsorption energy strength *E* (4*k*_*B*_*T*, 5*k*_*B*_*T*, 6*k*_*B*_*T* and 8*k*_*B*_*T*, corresponding to the absorption energy density from 6*k*_*B*_*Tnm*^−1^ to 11*k*_*B*_*Tnm*^−1^) under stretching force *f* = 0.3*pN* (FIG 3(b)). The interactions from weak to strong correspond to the salt concentrations from high to low. For appropriate absorption energy strengths (*E* = 4*k*_*B*_*T* and 5*k*_*B*_*T*, corresponding to absorption energy density from 6*k*_*B*_*Tnm*^−1^ to 8*k*_*B*_*Tnm*^−1^), the mean first passage time (MFPT) of nucleosome-wrapping is insensitive to the DNA supercoil density. Notably, under conditions of strong absorption (*E* ≥ 6*k*_*B*_*T*), the presence of negative supercoils serves to augment the efficiency of nucleosome wrapping.

**Fig 3.**
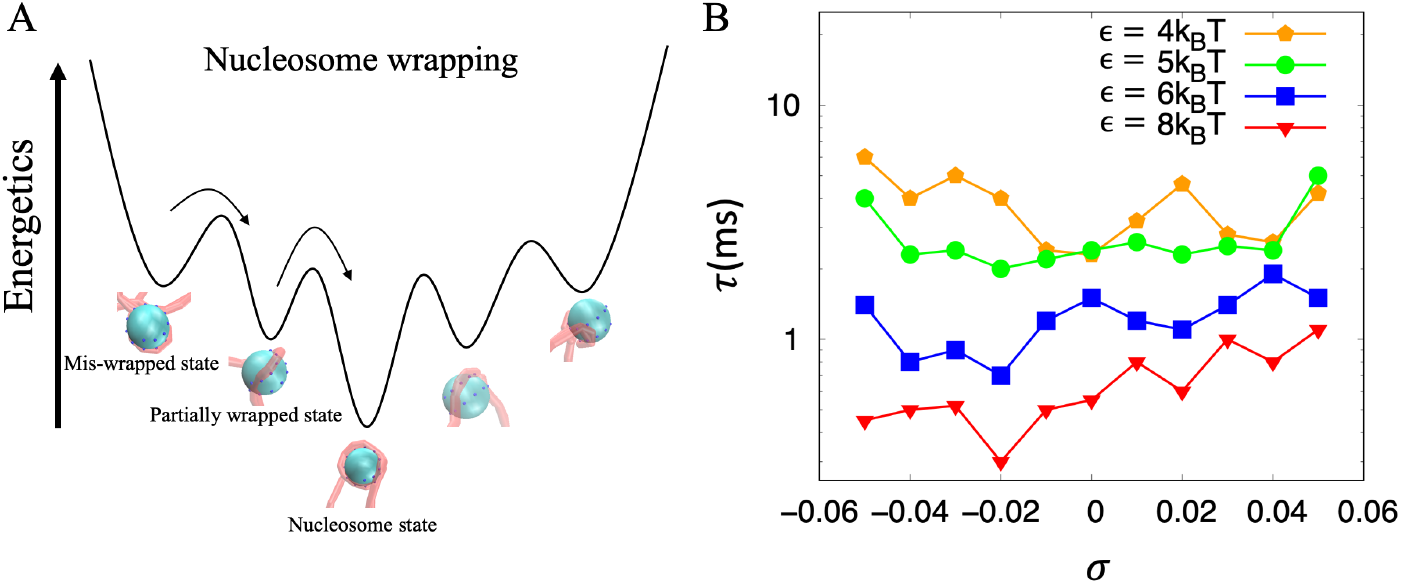
(A) Schematic illustration of the nucleosome wrapping process. Before completely nucleosome wrapping (Energetic minimum), the system explores a vast number of conformations of the mis-wrapped state (out of left-hand helix path) and partially wrapped state. (B) The mean first passage time (MFPT) for the completion of the nucleosome wrapping depends on the interaction energy strength *E* and the supercoil density *σ*. Each dot was obtained from 20 simulations trajectories with 10*ms* of each. For each given linking number, the DNA is stretched under constant force *f* = 0.3*pN*.

### Nucleosome-wrapping prefers negative supercoils

During transcription, the positive supercoils generated by the polymerases could dissociate the nucleosomes, while the negative supercoils behind might facilitate the re-assembly of the nucleosomes [9, 10]. Nevertheless, we have demonstrated that the nucleosome-wrapping process shows no sensitivity to the DNA supercoil density for the DNA-histone complex with an appropriate absorption energy strength under a small force (FIG.3b). The efficiency of nucleosome-wrapping is supposed to be force-dependent. Hence, we have compared the mean first passage time (MFPT) of nucleosome wrapping for supercoiled DNAs under different forces. (FIG 4(b)).

**Fig 4.**
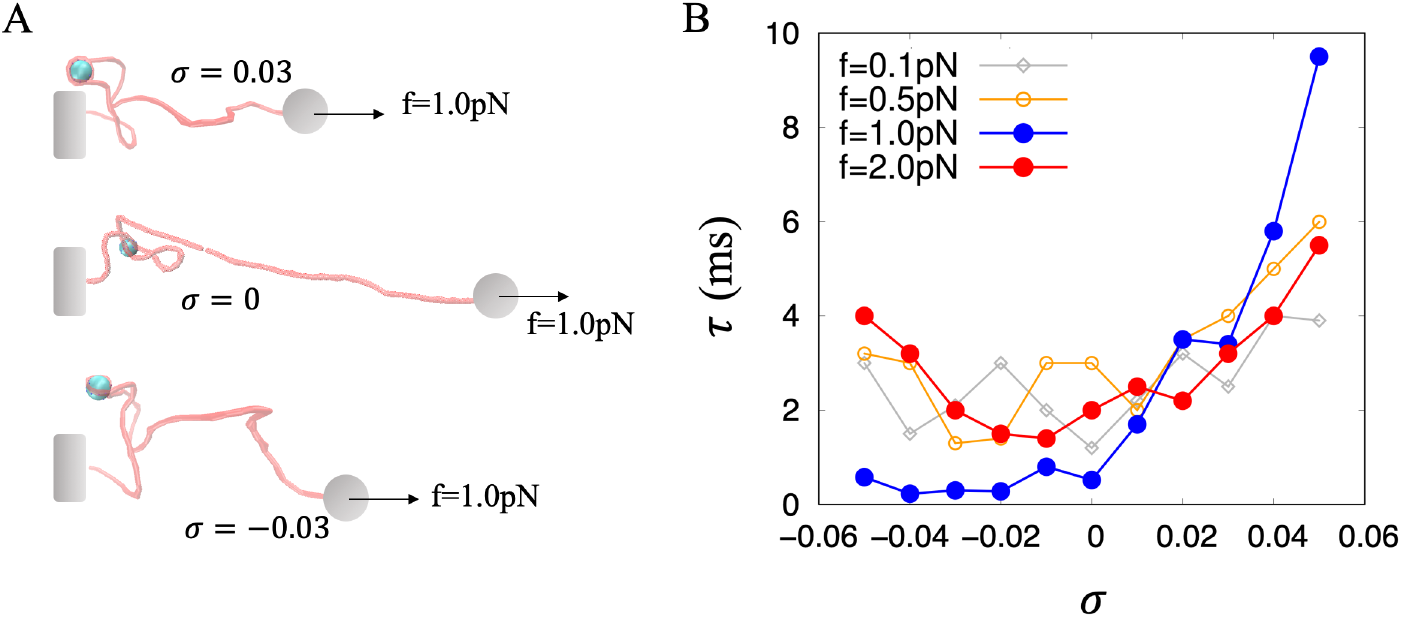
(A) Schematics for a single histone interacting with a supercoiled DNA under stretching force *f* = 1.0*pN*. (B) The supercoil-dependency of the MFPT for completing nucleosome wrapping. For the tensional DNA (*f* ≥ 0.5*pN*), nucleosome wrapping prefers the negative supercoiled DNA.

In the case of small applied force *f* = 0.1*pN*, the nucleosome-wrapping efficiency is insensitive to supercoil density. However, for tensional DNA, e.g, *f* ≥ 0.5*pN*, the MFPT for nucleosome wrapping by negative supercoiled DNA is significantly shorter compared to that achieved by positively supercoiled DNA. Notably, for the case of *f* = 1.0*pN*, the efficiency of nucleosome wrapping under negative supercoils is over an order of magnitude higher than that under positive supercoils.

Our simulations suggest that the DNA is interwound by the histone at forces *f* = 1.0 and 2.0*pN*, resulting in a relative extension 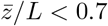. The change in DNA extension due to nucleosome-wrapping, 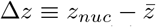 depends on the supercoiling (see FIG 5b). For the cases of *f* = 1.0 and 2.0*pN*, Δ*z*^(+)^ < Δ*z*^(−)^, leading to the work done being −*f* Δ*z*^(+)^ > −*f* Δ*z*^(−)^. The smaller work done by force *f* results in reduced energy barrier for nucleosome wrapping. Therefore, nucleosome wrapping tends to occur preferentially no negatively supercoiled DNA.

**Fig 5.**
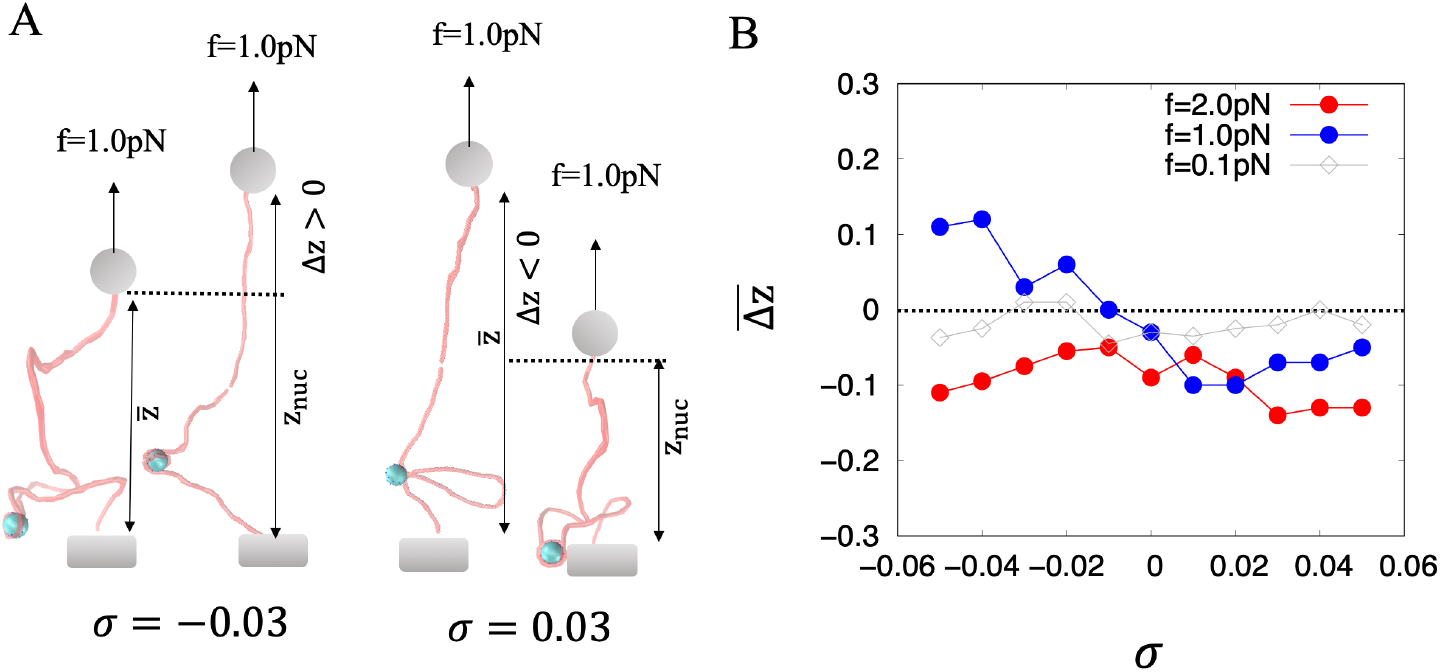
(A) The extension increment of the supercoiled DNA due to the nucleosome wrapping.Under *f* = 1.0*pN*, the nucleosome wrapping increases the extension of the negatively supercoiled DNA while reducing the extension of the positively supercoiled DNA. (B) The mean extension increment resulting from nucleosome wrapping is a function of supercoil density and stretching force. The positive increment in DNA extension due to nucleosome wrapping causes the decrease of the potential energy of the extended DNA, which is energetically favored.

## Discussion and Conclusion

For extendedly supercoiled DNA under *f* = 0.1, 1.0 or 2.0*pN*, the induced ΔΔ*Lk* = +1 on the unconstrained DNA is primarily attributed to writhe, with Δ*Wr* ≈ +1 and ΔΔ*Tw* ≈ 0 (FIG.6(a)). The increase in writhe rather than the change in twisting energy governs the dynamics. In other words, for positive writhe, |*Wr*| increases due to nucleosome-wrapping. In contrast, for negative writhe, |*Wr*| decreases due to nucleosome wrapping. Writhe measures the extent of DNA interwinding. Consequently, a significant magnitude of |*Wr*| is associated with a reduced Δ*z*. The negative supercoiling results in comparatively lower amount of work-done by *f*, implying a decreased energy barrier for nucleosome wrapping.

**Fig 6.**
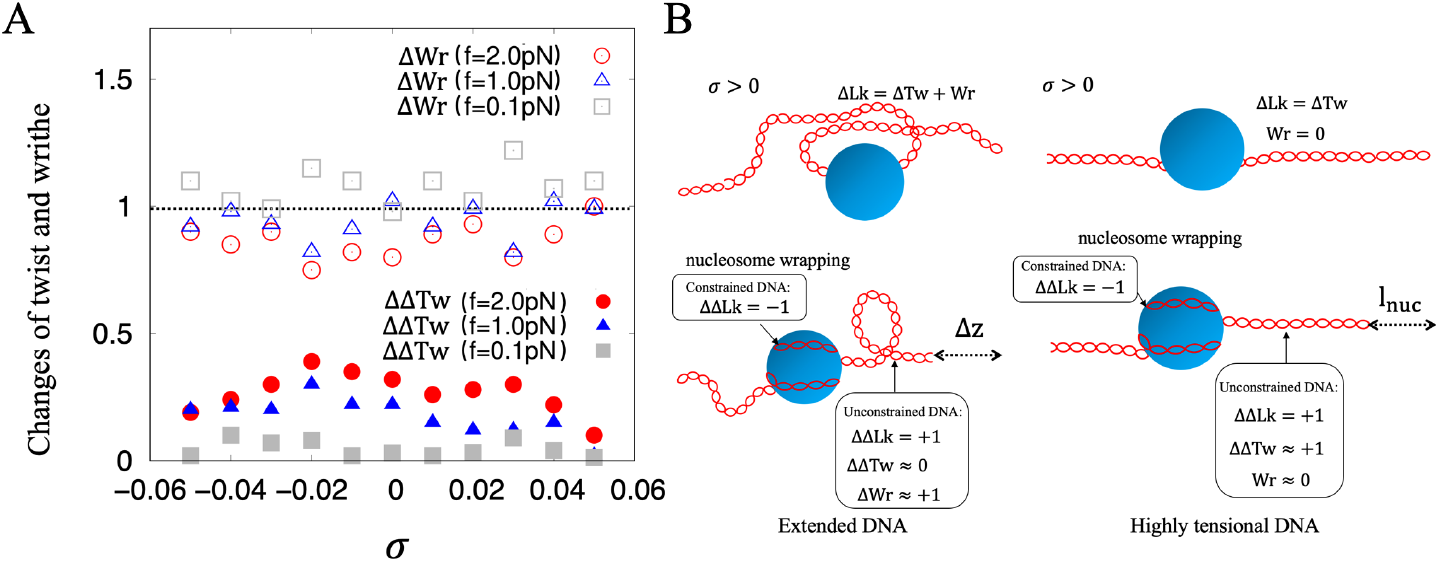
(A) Changes in the twist and writhe of the unconstrained DNA due to nucleosome-wrapping. (B)The changes in twist and writhe of (positive supercoils) DNA during winding around histone core. (Left) Extendedly (+) supercoiled DNA wraps around a histone under small force, e.g, *f* = 0.5, 1.0 and 2.0*pN*. The induced +1 linking number is mainly attributed to writhe, which causes work done during nucleosome wrapping. (Right) Highly tensional (+) supercoiled DNA wraps around a histone under large force, e.g, *f* = 10*pN*. The induced +1 linking number is mainly attributed to twist, which increases twisting energy during nucleosome wrapping.

In FIG.6(a), we observed that as the force *f* increases, there is a corresponding increase in in ΔΔ*Tw* and a decease in Δ*Wr*. For highly tensional DNA, such as those subjected to *f* = 5, 10 and 20*pN*, which are commonly utilized for stretching chromatin [12, 14], however, the increment of twist ΔΔ*Tw* ≈ 1 meanwhile writhe Δ*Wr* ≈ 0 during nucleosome forming, giving a rise to a supercoiling-independent change in extension, Δ*z* = *l*_*nuc*_. The change of the twisting energy is supercoiling-dependent, i.e., 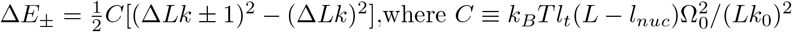 is the stiffness of the whole unconstrained DNA [36]. Evidently, Δ*E*_+_ > 0 > Δ*E*_−_, meaning nucleosome-wrapping always prefers negative supercoils.

In both of the cases, the energy barrier associated with nucleosome wrapping under positively supercoiled DNA exceeds that of negatively supercoiled DNA, thereby the preference for negative supercoiling.

In eukaryotic chromatin, DNA attains a highly compact structure due to its wrapping around histone cores, contributing to negative supercoiling. To elucidate the mechanism underlying the spontaneous wrapping of supercoiled DNA around histone proteins, we designed a Brownian dynamics model of the DNA-histone complex. We demonstrated that both the positively and negatively supercoiled DNAs can form the nucleosome conformation at indiscriminate time scales with appropriate absorption energies between DNA and histone. Furthermore, we observed that the preference for negative supercoiling during nucleosome wrapping becomes evident with increasing force. Several improvements to our model are warranted for future studies. The first one is the over-twist on the constrained DNA that wraps around histone, which may mainly originates from interaction with the histone rather than DNA intrinsic property [39]. Secondly, the nucleosome filament requires histone H1, which associates with the entry and exit of DNA from the histone core [21, 24]. The incorporation of the effects of the histone H1 allows nucleosomes to be arranged into a compact and stabilized array. In the large force regime, the dissociation of histone from DNA is dominating over re-assembling of nucleosome. Consequently, the net nucleosomes reassembling rate is determined by the difference between re-assembling rate and dissociate rate. Thus, the force-dependent kinetics of the nucleosome dissociation and re-assembling presents a promising research topic in the future.

## Supporting information

**S1 Text**. Dynamics of the rotational degrees

**S2 Text**. Pathway of nucleosome wrapping and the intermediate states

## acknowledgement

C.L. is supported by NSFC Grant #12005029 and start-up fund from Chongqing University of Posts and Telecommunication(A2020-029). B.W. is supported by NSFC Grant #12304246 and start-up fund from Wenzhou Institute, University of Chinese Academy of Sciences.

